# *Toxoplasma* induced cytokine release syndrome is critically dependent on the expression of pore-forming Perforin-Like Protein-1

**DOI:** 10.1101/2025.03.17.643671

**Authors:** Beth Gregg, Alfredo J. Guerra, Stephen A. Raverty, Aline Sardinha-Silva, Bjorn F.C. Kafsack, Tracey L. Schultz, Stephen J. Gurczynski, Bethany B. Moore, Vern B. Carruthers, Michael E. Grigg

## Abstract

Acute virulence in *Toxoplasma gondii* is linked to an excessive proinflammatory cytokine cascade during laboratory murine infection. Previous work showed that *T. gondii* secretes a pore forming protein, PLP1, that is required for efficient cytolytic egress from host cells. Deletion of the *PLP1* gene results in defective egress from infected culture cells and a marked reduction in parasite virulence. The goal of the present study was to gain insight into the nature of the attenuated virulence observed in *PLP1* knockout compared to wild type (WT) RH parasites. Using *in vivo* bioluminescence imaging, we show that parasites lacking PLP1 establish an acute infection and disseminate throughout the infected mice. Histological tissue analysis indicates that parasites cause severe pathology, even in the absence of PLP1. However, mice infected with *Δplp1* parasites evoke a protective inflammatory response, demonstrated by mouse survival and control of infection. Flow cytometric analysis was used to determine cellular changes occurring during both WT and Δ*plp1* parasite infection. Parasite control in the *Δplp1* infection was associated with earlier activation of myeloid cells and a moderate neutrophil response that, by comparison, becomes the dominant infiltrating cell type of WT infection. Positive disease outcome during *Δplp1* parasite infection is also associated with regulated induction of proinflammatory cytokines, including IFN-γ and TNF-α, and an earlier IL-10 regulatory response that is dysregulated during WT infection. Together these findings suggest a key role for *Toxoplasma* PLP1 in promoting a lethal inflammatory immune response during acute infection with a virulent strain of the parasite.

**Author Summary:** Pore-forming proteins are virulence determinants expressed by multiple different pathogens, with varied roles including cellular invasion and escape, immune cell destruction, and the hijack of host cell defenses. The pathogen, *Toxoplasma gondii*, expresses a pore-forming protein PLP1, that is required for cell lysis and acute virulence in mice. Here, we investigate the potential mechanisms by which this pore-forming protein promotes parasite virulence; from parasite replication and dissemination to immunologic outcomes after infection. *In vivo* infections demonstrate that parasites replicate, disseminate, and stimulate a protective immune response when PLP1 is not expressed. We show that PLP1 expression induces a parasite driven dysregulation of cell populations and cytokine/chemokine responses, resulting in cytokine release syndrome. In a broader context, *Toxoplasma’s* PLP1 is comparable to other pathogen pore-forming proteins that function as virulence determinants by their ability to alter host immune responses.

## Introduction

*Toxoplasma gondii* is an obligate intracellular parasite that can infect a wide range of intermediate hosts, including humans. In immunocompetent individuals, parasite infection usually results in mild symptoms (1). Generally, the host immune system mounts a strong interferon-gamma (IFN-γ)-dependent response which effectively controls the rapidly replicating tachyzoite stage during the acute phase of infection (2). During this acute stage, tachyzoites disseminate throughout the host, infecting multiple organs, including striated muscle and brain (3). Once the host mounts an effective immune response, *T. gondii* parasites differentiate into slowly replicating bradyzoites contained within a cyst that can persist for the life of the host (4). If the host becomes immunocompromised the bradyzoites may re-convert to tachyzoites causing various clinical manifestations of toxoplasmosis that can be fatal if untreated (5, 6). Acute toxoplasmosis during pregnancy can lead to transplacental dissemination and congenital toxoplasmosis resulting in developmental malformations, abortion, and even stillbirth (7).

North American strains of *T. gondii* consist mainly of three distinct clonal lineages: Type I, II, and III (8, 9), although other lineages, such as Type X, have also been identified (10–14). Type I strains are highly virulent in mice, with a lethal dose of a single parasite (9, 15). Conversely, Type II and III strains are much less virulent in mice with an LD_50_ of 1,000 or more parasites (15). In either case, a robust Th1 or type 1 immune response is necessary to induce protective immunity against *T. gondii*; however, infection may lead to overproduction of proinflammatory cytokines and death (16, 17). Prior work has highlighted the importance of IFN-γ in the control of both the acute and chronic stages of *T. gondii* infection (18–20). Early in the course of infection, interleukin-12 (IL-12) is released leading to an increase in the production of IFN-γ by T cells and NK cells (21–23). Additionally, tumor necrosis factor alpha (TNF-α) plays an important role in resistance to acute infection (24, 25). The difference in acute mortality between Type I and Type II/III strains has been attributed to an exaggerated host immune response (16, 17).

The *T. gondii* genome contains a member of the membrane attack complex/perforin (MACPF) protein family that is crucial for efficient cellular egress (26, 27). Deletion of the *PLP1* gene results in a striking reduction in parasite virulence (27). Despite the egress defect, *in vitro* replication of PLP1 deficient parasites was similar to that of the wild type (WT) RH strain (a highly virulent Type I strain). Whether the egress defect affects parasite expansion *in vivo* or if there is a change in the type and kinetics of the induced immune and cytokine response has not been systematically addressed in a mouse model. Our current structural and molecular understanding of the PLP1 mechanism of action (26, 28, 29) does not provide sufficient perspective to account for the virulence reduction shown by *PLP1* knockout parasites. In this study, we compared acute infection and host immune responses upon challenge with either WT or *PLP1* knockout (*Δplp1)* parasites. The results recapitulate our previous observation that the *PLP1* knockout parasites are markedly less virulent than the parent WT strain. To determine PLP1’s role during *in vivo* infection, we assessed whether the observed *in vitro* egress defect of *Δplp1* parasites contributed to a failure of the parasites to replicate and establish an acute infection *in vivo.* Specifically, we examined whether *PLP1* knockout parasites 1) failed to disseminate throughout the host, or whether 2) they caused less overall damage at the site of infection and/or 3) induced a protective immune response that regulated parasite proliferation. Our results showed that both *Δplp1* and WT parasites established an acute infection with parasite replication and multisystemic dissemination. However, in a dose-dependent manner, intraperitoneal inoculation of *Δplp1* parasites was associated with differences in the kinetics and absolute levels of immune cells and their cytokines and chemokines, histologic resolution of disease pathology coincident with the induction of a protective inflammatory response that regulated parasite infection. Our dataset offers new insight into the mechanism of PLP1 action and its contributing role in acute virulence.

## Results

### RH and *Δplp1* parasites both establish acute infections

Prior work showed that *T. gondii* parasites deficient in PLP1 were avirulent in outbred mice (27). Using a C57BL/6 inbred mouse model, mice were infected either with WT (RH) or *Δplp1* parasites. As expected, WT-infected mice succumbed to acute toxoplasmosis between days 8 and 11 post-infection (dpi), whereas all doses of *Δplp1* infected mice survived infection (**Fig 1A**). One interpretation for this observation is that *Δplp1* parasites failed to establish an acute infection. To assess this prospect we infected mice intraperitoneally with 10^4^, 10^5^, and 10^6^ *Δplp1* or 100 WT RH tachyzoites and monitored the infection by bioluminescence imaging (**Fig 1**). At the lowest inoculation, *Δplp1* infection levels were similar to that of WT up to 4 days post-infection (**Fig 1B**). Compared to WT, high doses of 10^5^ and 10^6^ *Δplp1* parasites exhibited significantly higher parasite burdens early in infection through to day 5/6 post-infection. By 6 dpi parasite loads were equivalent between WT and *Δplp1*, after which *Δplp1* parasites were eventually controlled and loads decreased 2-3 logs by day 11 (**Fig 1C and D**). In contrast, WT parasite load continued to increase after day 6 until mice succumbed to infection between 8-11 dpi. Importantly, at all infectious doses, PLP1 was not required for parasites to establish acute infection and to proliferate *in vivo* (**Fig 1B – D**). However, PLP1 or some other factor(s) dependent on PLP1 function, was required for WT parasites to continue to proliferate to unrestricted levels and be acutely virulent.

**Fig 1.**
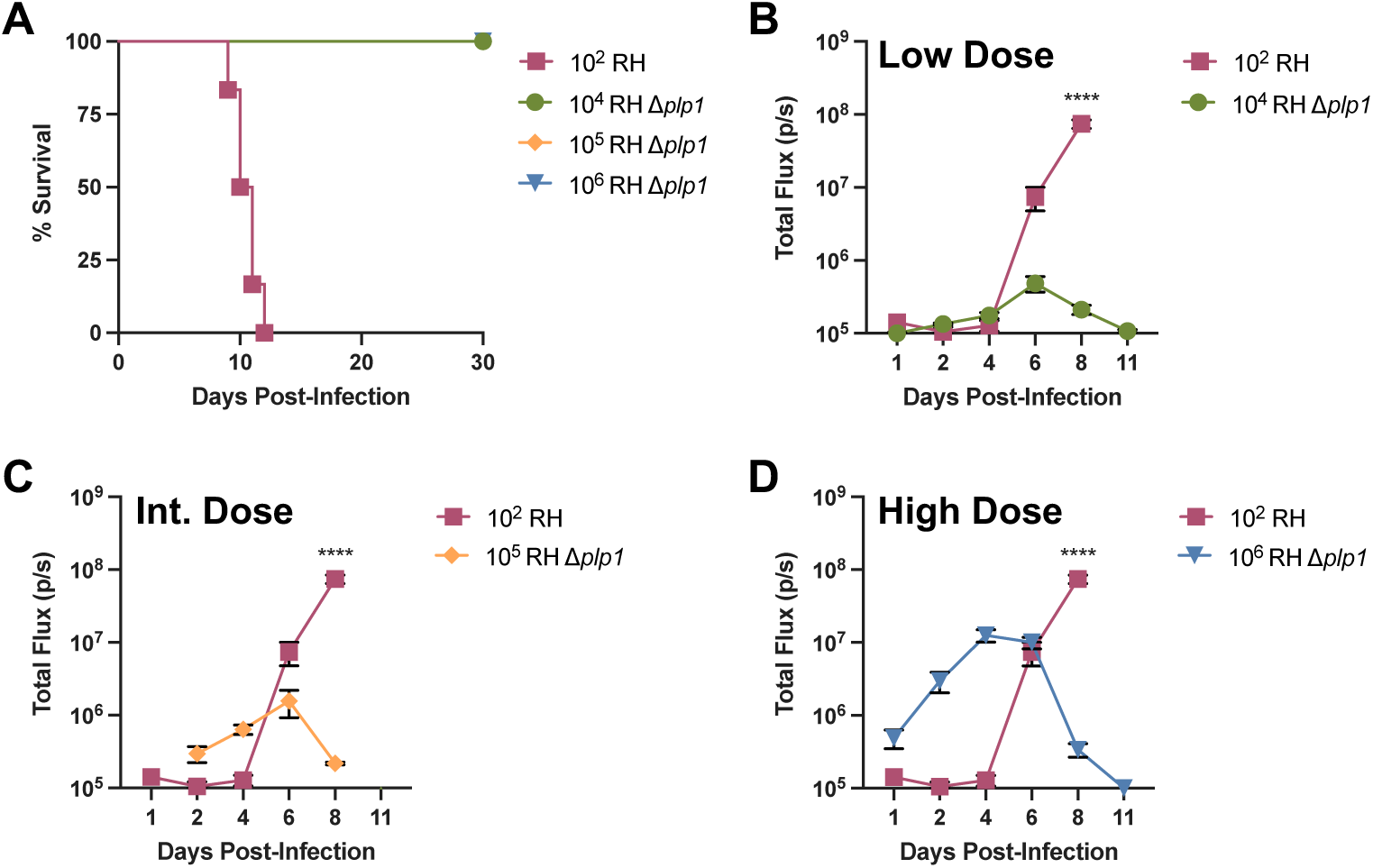
PLP1 deficient parasites establish acute infection. (A) C57BL/6 mice intraperitoneally (i.p.) infected with 10^2^ WT (RH), or low dose 10^4^, intermediate (int.) dose 10^5^, or high dose 10^6^ RH *Δplp1* parasites were monitored for survival. Mice infected with wild type parasites succumbed to the infection between days 9 and 12 post-infection. Parasite burden was monitored by bioluminescence imaging of parasite expressed firefly luciferase after intraperitoneal infection with 10^2^ WT, 10^4^ (B), 10^5^ (C), or 10^6^ (D) tachyzoites. Data are pooled from 2 independent experiments with 6 total mice. Error bars indicate standard error mean (SEM). **** is equivalent to p<0.0001, in a two-way ANOVA.

### RH and *Δplp1* parasites disseminate throughout infected mice

To investigate whether the reduced virulence in Δ*plp1* parasites was due to an intrinsic inability of the mutant parasites to disseminate throughout the host during the course of the infection, mice were infected intraperitoneally with 10^2^ WT versus 10^4^ (low dose) or 10^6^ (high dose) *Δplp1* luciferase-expressing tachyzoites and sacrificed at 4, 6, 8, and 11 dpi. During experimental infection with *T. gondii*, Type I (RH) strains rapidly disseminate to a wide variety of tissues (16, 17). For these studies, we chose to inoculate mice with both the low dose (10^4^) and the high dose (10^6^) of tachyzoites because these doses had similar kinetics and progression of infection compared to the 10^2^ dose of WT parasites at either early or later time points, respectively (**Fig 1B-C**). Similar to whole body imaging of infected mice, BLI of excised internal tissues confirmed that both doses established infection. Infection with 10^4^ *Δplp1* and WT parasites demonstrated tachyzoite replication primarily within the peritoneal cavity until 4 dpi, after which parasites could be detected in peritoneal proximal tissues of the intestine, liver, and spleen (**Fig 2F, G, H, and D**). In contrast, the 10^6^ *Δplp1* parasites disseminated a few days earlier from the initial site of injection and achieved higher parasite loads in all tissues examined (peritoneum, lung, heart, liver, spleen, intestine) at early time points. Importantly, by 6 dpi, parasite loads in the WT infected mice caught up and were essentially the same as the 10^6^ *Δplp1* dose, measured using an organ specific ROI (**Fig 2B-H)**. By 8 dpi, parasite burden in both doses of *Δplp1* infection had peaked or was already decreasing in all tissues, whereas WT parasites continued to expand and were not controlled. Overall, these results suggest that PLP1 is not required for *Toxoplasma’s* ability to disseminate, but that PLP1, or some other factor(s) associated with PLP1 function, is virulence enhancing and promotes unrestricted growth of parasites in mice until they succumb to acute infection.

**Fig 2.**
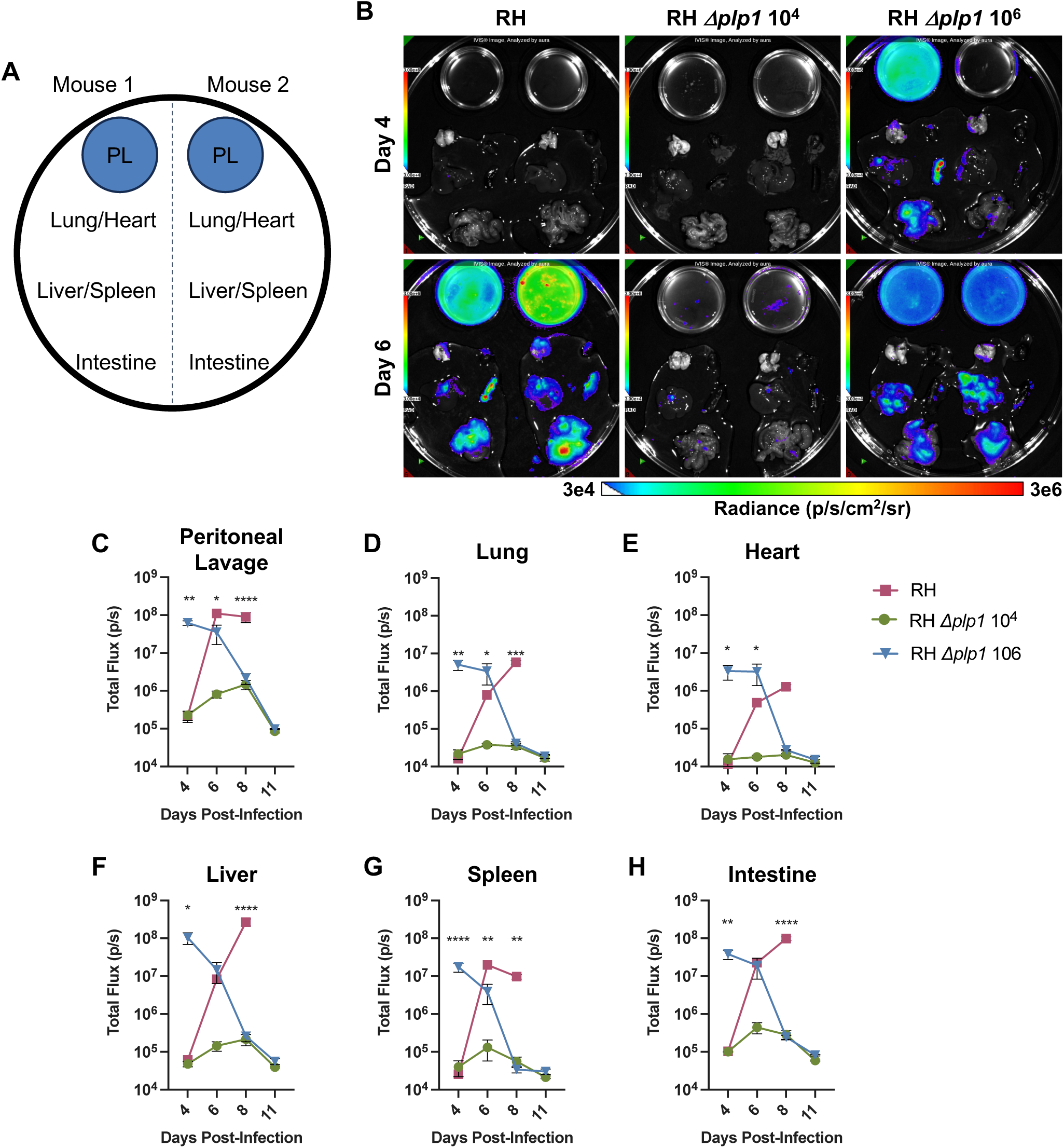
PLP1 deficient parasites disseminate to distal tissues during acute infection. Schematic of tissue placement for bioluminescence imaging (A). Representative images of tissues on days 4 or 6 post-infection, from C57BL/6 mice infected i.p. with either 10^2^ WT (RH, left), 10^4^ Δ*plp1* (middle), or 10^6^ RH Δ*plp1* (right). Scale represents radiance from 3e4 (blue) to 3e6 (red) photons/s/cm^2^/sum of radiance. Calculation of bioluminescence in regions of interest delineating each tissue, to measure parasite dissemination throughout the organism: peritoneal lavage (C), lung (D), heart (E), liver (F), spleen (G), and intestine (H). Representative results from two independent experiments. Error bars indicate SEM and statistical analysis was performed by two-way ANOVA (n=3, * p<0.05, ** p<0.005, *** p<0.0005, **** p<0.0001).

### Histopathology and parasite burden

We next investigated the histopathology associated with infection at 6 and 8 dpi, time points when parasite load diverged between WT (unrestricted growth) and *Δplp1* (controlled growth) parasites. Qualitative assessment of sectioned tissues from all challenge cohorts (10^2^ WT, 10^4^ or 10^6^ *Δplp1* doses) identified among the different treatment groups similar pathologic lesions that varied in both their frequency and extent of inflammation and necrosis. The most consistent histopathologic feature observed was the detection of intralesional protozoa that was dependent on infection inoculum. Histologically, parasite loads and distribution throughout tissues were similar in both WT RH and *Δplp1* 10^6^ doses at both days 6 and 8, whereas the low dose of *Δplp1* (10^4^) had fewer parasites at day 6 and none observed at day 8 (**Fig 3A**). Within the spectrum of lesions, more areas of extreme necrosis and inflammation were consistently found in the *Δplp1* high dose (10^6^) infection at both 6 and 8 dpi (**Fig 3**).

**Fig 3.**
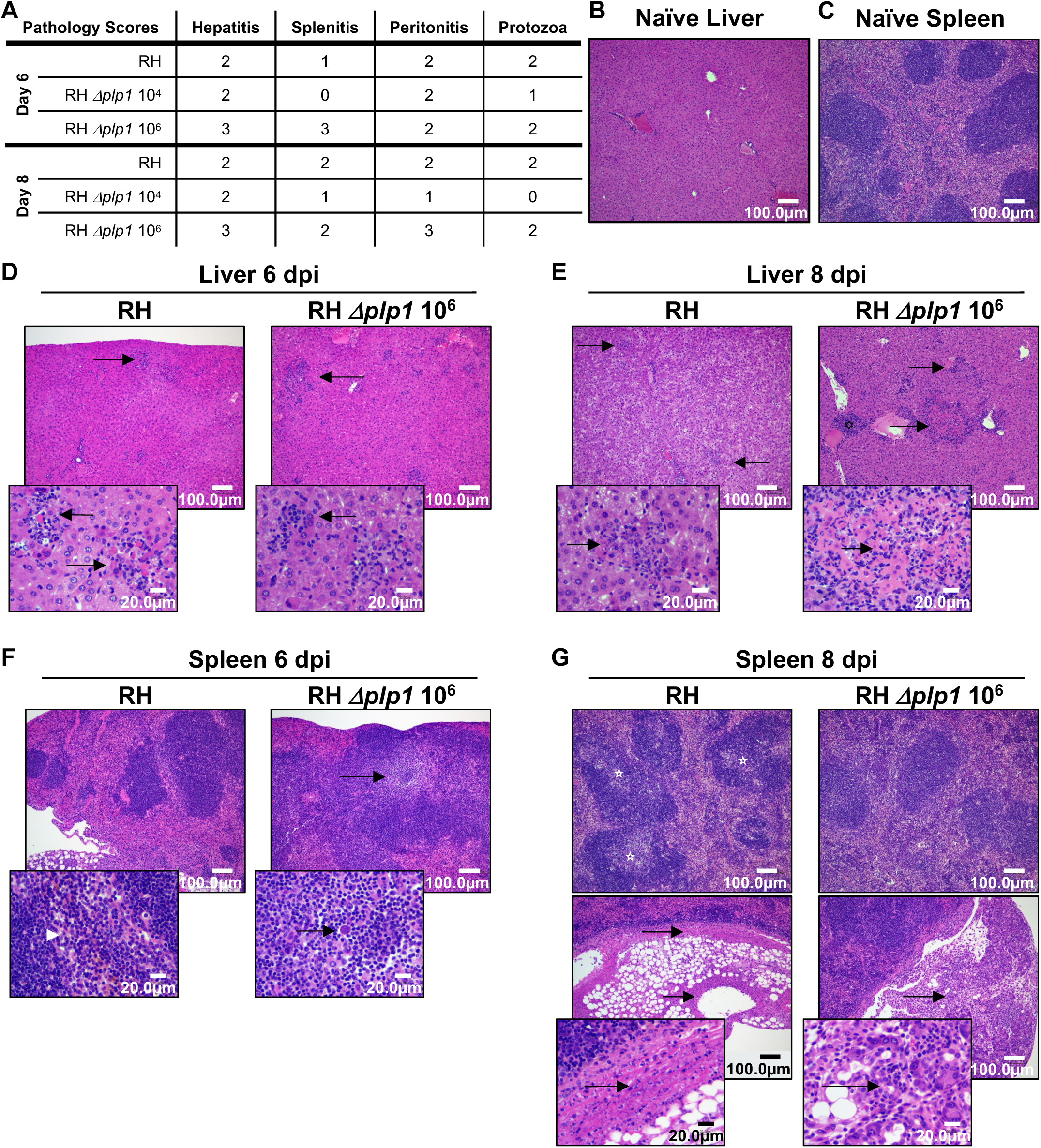
PLP1 deficient parasites induce similar pathology to wild-type RH controls. H&E stained tissue from WT (RH 10^2^)and 10^6^ RH *Δplp1* infected mice were assessed for evidence of infection induced pathology on days 6 (D & F) and 8 post-infection (E & G), and compared with representative naïve liver (B) and spleen (C). Liver images (B, D, and E) were taken at 10x magnification with insets (60x magnification) to highlight regions of inflammation, necrosis and intralesional parasites. Sections of spleen (C, F, and G) were imaged at 4x magnification with 60x magnification insets to highlight granulopoiesis (F), peritonitis, lymphoid depletion (stars), and capsular splenitis (G). Average hepatitis, splenitis, peritonitis, and intralesional protozoa pathology scores for WT and 10^6^ *Δplp1* infected mice for 6 and 8 dpi are included in a table (A).

During the peak of infection, pyogranulomatous hepatitis, characterized by a predominant influx of neutrophils and macrophages, was consistently identified among the different infection groups. Additionally, there was hepatocellular dissociation, degeneration, apoptosis, and necrosis. Fibrinous capsular hepatitis was also observed (**Fig 3A**). In mock-infected mice, the liver sections had normal architecture with no evidence of underlying pathology, infection or inflammation (**Fig 3B**). The most significant histopathology observed in the livers of all infected mice was mild to moderate multifocal and random nonsuppurative hepatitis (**Fig 3A and D-E**). Of note, the inflammatory infiltrate and hepatocellular necrosis were more intense, and the lesions more disseminated throughout the parenchyma of the *Δplp1* high dose (10^6^) infection than WT at both days 6 and day 8 (**Fig 3D-E).**

Histopathology of the spleen featured mild and moderate multifocal splenitis with variable lymphoid hyperplasia and occasional depletion, fibrin deposition, and capsular splenitis in each cohort; however, the frequency, intensity, and extent of the lesions varied by treatment dose and post inoculation time within each challenge cohort (**Fig 3F, G**). Naive spleen showed no indication of underlying pathology or infection, with normal well demarcated white and red pulp tissue architecture (**Fig 3C**). At 6 dpi, the spleen from WT infected mice featured mild reactive change, characterized by lymphoid hyperplasia (**Fig 3A and F**). In WT infected mice specifically, there was evidence of moderate to marked multifocal splenitis apparent throughout the stroma, which extended along the splenic capsule, and occasionally spanned across to the serosal surface of adjoining fat lobules (peritonitis) and infiltrated deep into pancreatic lobules (pancreatitis) (**Fig 3F**). A myriad of protozoa were interspersed within the inflammatory exudate (**Fig 3F**). At higher magnification, areas of extramedullary myelopoiesis including immature and developing neutrophilic granulocytes, were more pronounced in the WT infected mice (**Fig 3F**). Fewer areas of cellular proliferation were observed in the spleens of the high dose (10^6^) *Δplp1* infected mice. This disparity suggests that extramedullary generation of immune cells (reviewed in (30, 31)), and the overall immune response, was significantly altered by parasite expression of PLP1. Day 6 spleens from *Δplp1* 10^6^ dose infected mice exemplified a more chronic (inflammation score 3) fibrinous peritonitis or fibrinous capsular splenitis that possessed numerous intralesional and extracellular protozoa tracking along the serousal surface and extending into the stroma (**Fig 3F**). By day 8, the splenic inflammation of *Δplp1* infected mice was even more extensive and extended into the stroma of the adipose tissue as well as the interstitium of the pancreas (**Fig 3G**). Notably, WT-infected mice, but not *Δplp1* infected mice, had a general pallor (pale appearance) within the central portions of the follicule (see starred follicules), and hypocellularity within the intervening white pulp. Lymphoid depletion was evident (**Fig 3G**). Potential explanations for the dramatic depletion of lymphocytes was either a peripheral mobilization of the lymphocytes to other areas of inflammation within the infected mice, reduced lymphopoiesis, or that it was the direct result of lymphocytolysis, as previously observed during *Toxoplasma* infection of mice (32).

### Infection with RH and *Δplp1* parasites results in differential myeloid cell recruitment

To gain insight into the host response to infection with *Δplp1* parasites we investigated the composition of immune cells that infiltrated the peritoneal cavity after i.p. inoculation. Mice infected with either WT or *Δplp1* (low vs. high doses) parasites initiated a steady increase in peritoneal exudate cells (PECs) that appeared to plateau between 6 and 8 dpi (**Fig 4A**). After 6 dpi and through 11 dpi, total PEC numbers were maintained at high levels in response to *Δplp1* parasites, whereas PEC numbers declined precipitously just prior to mouse death in WT infected mice (**Fig 4A**). Host cell recruitment to the peritoneal cavity was partially dependent on parasite load. Mice infected with 10^6^ *Δplp1* parasites had a significantly higher number of PECs than WT infected mice between 3 and 6 dpi, but approximately the same at 6 dpi when parasite load was equal (**Fig 4A and B**). Consistent with these results, an equivalent number of PECs were recruited through the first three days of WT versus 10^4^ *Δplp1* infection when parasite load was similar. Of note, PEC numbers remained equivalent at 6 dpi despite significantly different parasite burdens (**Fig 4B**). We next investigated whether resident peritoneal macrophages (CD11b^+^/F4/80^+^, gating strategy **S1 Fig**), which comprise part of the acute phase immune response, were differentially impacted during infection with either WT or *Δplp1* (10^4^ and 10^6^ doses). These cells are known to rapidly disappear during immune stimulation, the result of cell death or sequestration to other anatomic sites such as the omentum (33, 34). No difference was detected; all infection groups depleted F4/80^+^ macrophages by day 4 post-infection (**Fig 4C**). These results support the conclusion that *Δplp1* parasites stimulate the early host immune response in a manner similar to WT parasites.

**Fig 4.**
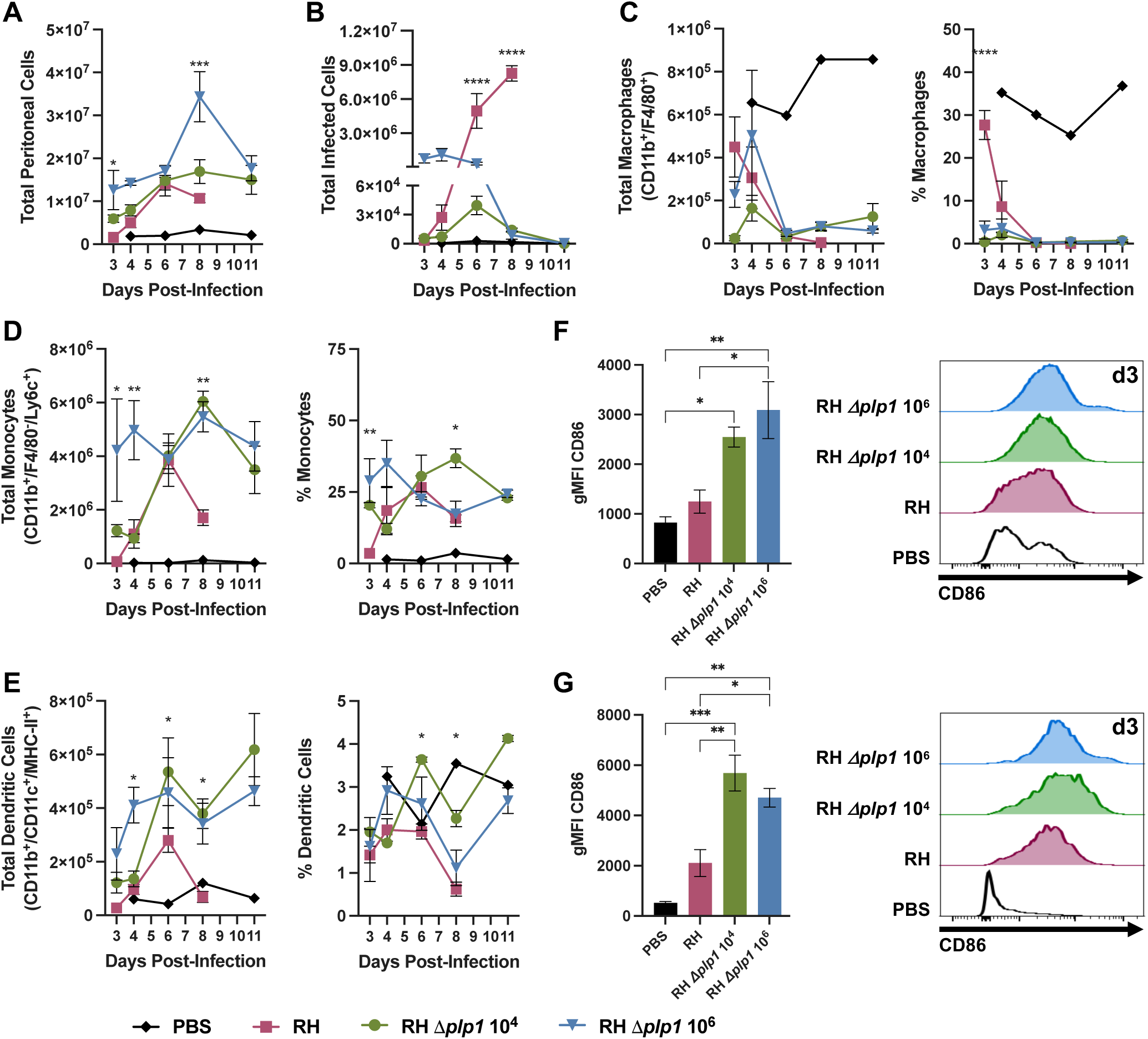
Peritoneal cells are rapidly depleted during infection with WT parasites, but not by PLP1 deficient parasites. C57BL/6 mice were infected via i.p. inoculation with either 10^2^ WT (RH, red), 10^4^ (green) or 10^6^ (blue) RH *Δplp1 Toxoplasma gondii*, and peritoneal cells were isolated and analyzed at days 3, 4, 6, 8, and 11 post-infection. Total numbers of peritoneal cells (A), GFP^+^ infected cells (B), CD45^+^/CD11b^+^/F4/80^+^ macrophages (C), CD45^+^/Ly6G^−^/CD11b^+^/F4/80^−^/Ly6C^+^ monocytes, and CD45^+^/Ly6G^−^/CD11b^+^/CD11c^+^/MHC-II^+^ dendritic cells (E), as well as their respective percent of the total cellular population are shown. Geometric mean fluorescence intensity (gMFI) of costimulatory CD86 on monocytes (F) and dendritic cell (G) populations was analyzed by flow cytometry and plotted or displayed as a representative histogram for day 3 post-infection (modal axis). Flow cytometry data are representative of two individual experiments with n=3 or 4 animals. PBS controls (black) are from one animal on each represented day post-infection. Included statistical analyses were performed by either one-way (CD86 expression) or two-way ANOVA; p values represent significance as compared with RH (* p ≤ 0.05, ** p ≤ 0.005, *** p ≤ 0.0005, **** p ≤ 0.0001). All RH infected mice succumbed at days 8 or 9 post-infection.

Next, we assessed the levels of both monocytes and dendritic cells in the PECs of mice infected with WT versus *Δplp1* parasites. These immune cells play key roles in controlling *T. gondii* infection and virulence by their ability to control parasite growth directly and/or activate T cell responses (35, 36). A significant difference in total monocytes, characterized as Ly6G^−^/CD11b^+^/F4/80^−^/Ly6C^+^, and dendritic cells (Ly6G^−^/CD11b^+^/CD11c^+^/MHC-II^+^) was detected early during infection (between 3-6 dpi) that was dependent on parasite load (**Fig 4D and E**). By 6 dpi, all infection groups had nearly equivalent numbers of Ly6C^+^ monocytes, but the RH infection group demonstrated a trend toward less CD11c^+^ dendritic cell recruitment (**Fig 4D and E**). As observed with total PECs and by histopathology, WT but not *Δplp1* infected mice experienced a sharp, highly significant decline in both monocyte and dendritic cell populations by 8 dpi, just prior to mouse death (**Fig 4 D and E**). Both *Δplp1* infection groups maintained high levels of monocytes and dendritic cells through 11 dpi as mice recovered from acute infection (**Fig 4 D and E**). Interestingly, diminished dendritic cell recruitment was previously described by Tait et al. 2010, when WT RH virulence was correlated with the impairment of dendritic cell recruitment to the PECs (36). In addition to the decrease in dendritic cells during WT RH infection, early activation of these cells was likewise impaired (36). To investigate whether the reduced virulence observed in *Δplp1* infected mice correlated with an increased activation of antigen presenting cells, mean fluorescence intensity of CD86 on both monocytes and dendritic cells was assessed by flow cytometry. At 3 dpi, CD86 expression was significantly increased on both monocytes and dendritic cells in *Δplp1* but not WT infected mice (**Fig 4F and G**). Taken together, both RH WT and *Δplp1* infection stimulate the host response, but Δ*plp1* parasites did not disrupt early PEC antigen presenting cell recruitment and activation compared to WT.

Finally, we measured the absolute level and percentage of neutrophils in the PECs (**Fig 5A**). Mice infected with WT parasites showed a statistically significant increase in both total neutrophils and frequency of neutrophils, compared to *Δplp1* infected mice on 6 and 8 dpi, as determined by two-way ANOVA (**Fig 5A**), which supported the histopathology findings. By 8 dpi, neutrophils were the predominant cell type found in the PECs of WT infected mice, reaching on average 59% of total cells. Despite the high numbers of neutrophils, WT parasite growth failed to be controlled during infection (**Fig 4B**). In contrast, while mice infected with *Δplp1* parasites maintained an elevated percentage of neutrophils (11% average) compared to naïve mice (2%), the total numbers and percent of PECs remained relatively constant throughout the infection time course (**Fig 5A**). While studies have determined that the role of neutrophils during acute infection with a Type II *T. gondii* strain is minimal, no studies have specifically investigated the potential for a pathologic role of neutrophils during a Type I infection (35, 37–39).

**Fig 5.**
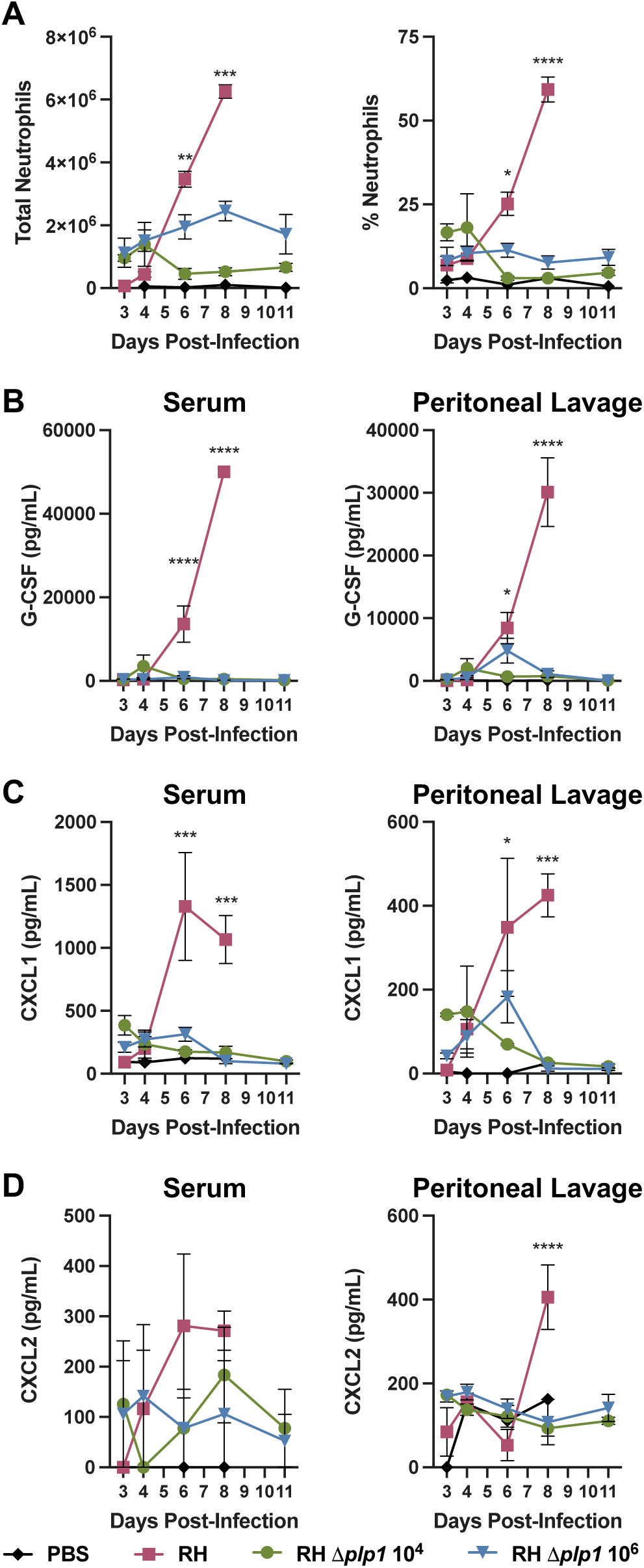
Dampened neutrophil and neutrophil chemoattractant response during *Δplp1* parasite relative to wild type infection. 10^2^ WT (RH, red), 10^4^ (green) RH *Δplp1*, or 10^6^ (blue) RH *Δplp1* were administered i.p. to C57BL/6 mice, and total or percent neutrophils (CD45^+^/CD11b^int^/Ly6G^+^) present in the peritoneal cavity was measured by flow cytometry on days 3, 4, 5, 6, and 11 post-infection (A). Concentration of neutrophil growth factor G-CSF (B), and chemoattractants KC/CXCL1 (C) and MIP2/CXCL2 (D) were measured by Milliplex analysis in both serum and 2 mL peritoneal lavage wash throughout infection. Data are representative results from of two individual experiments with n=3 and 4 mice per group/PBS control n=1 mouse per day. * represents p ≤ 0.05, ** p ≤ 0.005, *** p ≤ 0.0005, **** p ≤ 0.0001 **** represents p ≤ 0.001 as determined by two-way ANOVA. All WT infected mice succumbed to infection by days 8 and 9 post-infection.

### Host response to wild type vs. *Δplp1* has an altered chemokine response

To determine whether the PLP1-dependent increase in peritoneal cavity neutrophil abundance in WT infected mice was the result of increased production and recruitment, the temporal abundance of growth factor G-CSF and neutrophil chemoattractants CXCL1 and CXCL2 were investigated. As a key driver of neutrophil production, maturation, and activity, differences in G-CSF abundance could explain the observed increases in PEC neutrophil numbers during WT infection (40). Only WT infection resulted in significantly increased levels of G-CSF starting after 4dpi in both the serum and peritoneal lavage (Fig 5B, and S2 Fig). Furthermore, only WT infection resulted in significant increases in neutrophil chemoattractants. Specifically, all mice showed early systemic (3 and 4dpi serum) increases in CXCL1 and CXCL2 levels, although these values did not reach significance compared to naïve mice (**Fig 5C and D**). Local concentrations of CXCL1 in the peritoneal lavage fluids on 3 and 4dpi in all cohorts also did not reach statistical significance, but were elevated compared to uninfected mice (**Fig 5C**). In mice infected with *Δplp1* parasites, the CXCL1 levels plateaued and returned to pre-infection levels by 8 dpi, correlating well with total neutrophil numbers in the PECs (**Fig 5A and C**). However, in the WT infected mice, increased levels of both serum and peritoneal CXCL1/CXCL2 were statistically significant on 6 and 8dpi in a PLP1 expression-dependent manner (**Fig 5C and D**). Importantly, the increased levels of G-CSF and neutrophil chemoattractants could not be attributed solely to excessive parasite numbers, as the high dose *Δplp1* infection did not result in neutrophil and chemokine levels compared to those observed in the WT infection at its peak (**Fig 5**). These findings suggest that robust recruitment of neutrophils to the site of infection in WT infected mice is associated with fatal disease, whereas more tempered neutrophil recruitment during *Δplp1* infection is linked to survival.

### Host response to wild type vs. *Δplp1* has an altered cytokine response

In addition to changes in the myeloid compartment during infection, we reasoned that differences in cytokine responses to WT versus *Δplp1* infection might also provide insight into the basis for distinct survival outcomes. It is important to note that previous studies have shown that cytokine driven immunopathology and mouse survival is critically dependent on parasite burden (16, 17). A low dose (10^2^ tachyzoites) infection with Type II *T. gondii* is avirulent, whereas a high dose (10^5^ tachyzoites) infection leads to cytokine overproduction with all mice dying acutely, similar to a Type I RH infection (10^2^ tachyzoites) (17). For this reason, cytokine responses were assayed using both a low and high dose of *Δplp1* parasites and compared against WT infection at 10^2^ tachyzoites. Initially, Th1 cytokines known to be critical in priming immunity against *T. gondii* infection were investigated. IL-12p40 levels were significantly higher in WT infected mice compared to *Δplp1* infected mice, both locally by 6 dpi and systemically by 8 dpi (**Fig 6A and E**). The kinetics and absolute level of IFN-γ produced in WT infected mice peaked locally at 6 dpi before decreasing, and systemically at 8 dpi, just prior to mouse death (**Fig 6B and F**). Likewise, both doses of *Δplp1* parasites induced significant levels of systemic IFN-γ compared to naïve controls, but the levels were at least 4-fold lower compared to WT infection at 6 dpi (**Fig 6F and S2 Fig**). Local IFN-γ levels correlated with parasite load. Accordingly, IFN-γ was equivalent 6 dpi when WT and *Δplp1* (10^6^ dose) parasite burden was equal, whereas IFN-γ was significantly lower in the *Δplp1* (10^4^ dose) when parasite burden was lower (**Fig 6B**). By 8 dpi, IFN-γ dramatically decreased in *Δplp1* infected mice in concordance with the control of parasite replication in the peritoneal cavity, but remained elevated in WT infected mice where parasite burden continued to increase (**Fig 6B and F, and S2 Fig**). Responses observed for IL-18 and TNF-α (**Fig 6C, G and 6D, H**, respectively) demonstrated the same pattern and kinetics as the IFN-γ responses. Considering that a high dose *Δplp1* infection, which fail to cause acute mortality, induced peak local cytokine levels and parasite burden comparable to lethal WT RH or high dose Type II strain infections (17), our results suggest that PLP1 and/or factors associated with its expression, combined with parasite load, is sufficient to generate a lethal overstimulation of the immune response.

**Fig 6.**
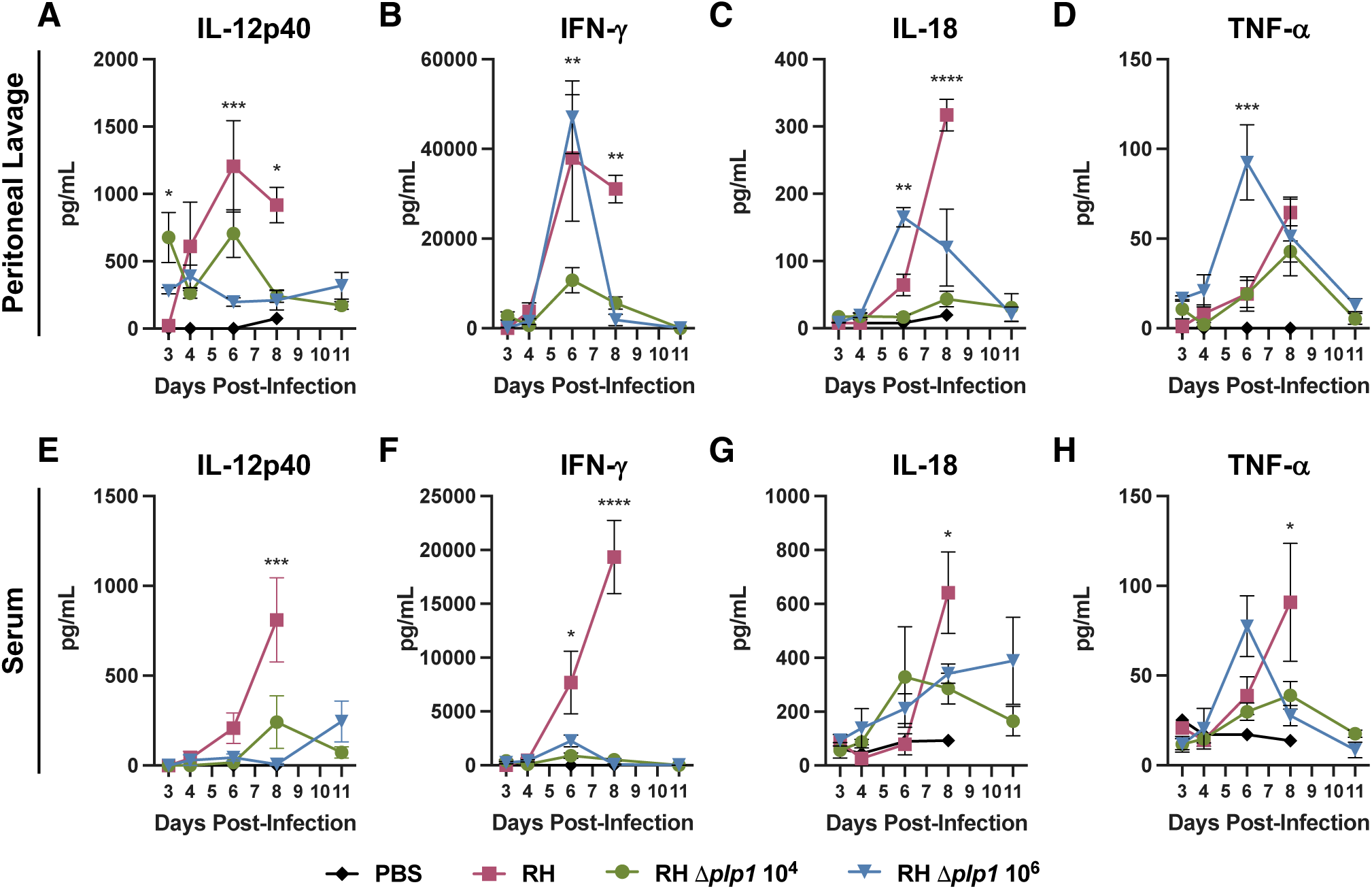
Altered cytokine responses in mice infected with PLP1 deficient parasites relative to those infected with wild type parasites. Levels of IL-12p40 (A, E), IFN-γ (B, F), IL-18 (C, G), and TNF-α (D, H) in the PEC lavage (A-D) and serum (E-H) were assessed by cytokine ELISA and Milliplex on days 3, 4, 6, 8 and 11 post-infection. C57BL/6 mice were infected i.p. as previously described. Data are representative of two individual experiments with WT (RH 10^2^ tachyzoites, red), low dose RH *Δplp1* (10^4^ tachyzoites, green), and high dose RH *Δplp1* (10^6^ tachyzoites, blue). PBS injected control mice (black) were included across the infection time course. Experiments were analyzed by two way ANOVA (* p ≤ 0.05, ** p ≤ 0.005, *** p ≤ 0.0005, **** p ≤ 0.0001 ** represents p ≤ 0.01, *** represents p ≤ 0.001).

Previous studies have demonstrated that upregulation of regulatory cytokines (such as IL-10) often peak after a robust Th1 proinflammatory response is generated by Type I RH parasites, but that this precipitous expansion of regulatory cytokines is insufficient to control dysregulated immune pathology and mouse death (17). This is because lethal doses of *Toxoplasma* typically generate a “cytokine storm” characterized by a failure to regulate the overproduction of IFN-γ, TNF-α, IL-18, and nitric oxide (17, 41, 42). Accordingly, mice infected with WT parasites showed significant peritoneal IL-10 levels 8 dpi; two days after IFN-γ levels peaked and just prior to succumbing to the infection (**S1A and G**). However, in mice infected with *Δplp1* parasites, IL-10 increased only during the high dose infection and was concomitant with the IFN-γ response, consistent with homeostatic regulation of a localized proinflammatory state (**Fig 7A and G**). In addition to hyberbolic IL-10, other hallmarks of cytokine release syndrome include the induction of dysregulated levels of IL-6, CCL2, CCL3, and CCL4 (43). Investigation into the concentrations of these cytokines, as well as CCL11 and G-CSF, during WT infection demonstrated significant increases both locally and systemically, whereas *Δplp1* parasites induced relatively small, if any, increases that resolved by 11 dpi (**Fig 5B**, **Fig 7B-F, 7H-L, and S2 Fig**). Taken together our findings suggest that while the production and kinetics of Th1 cytokines, including IFN-γ, were similar in mice infected with WT and high dose *Δplp1* parasites, survival of *Δplp1* infected mice was associated with an appropriate regulation of cytokines and chemokines that precluded the generation of a systemic cytokine release syndrome.

**Fig 7.**
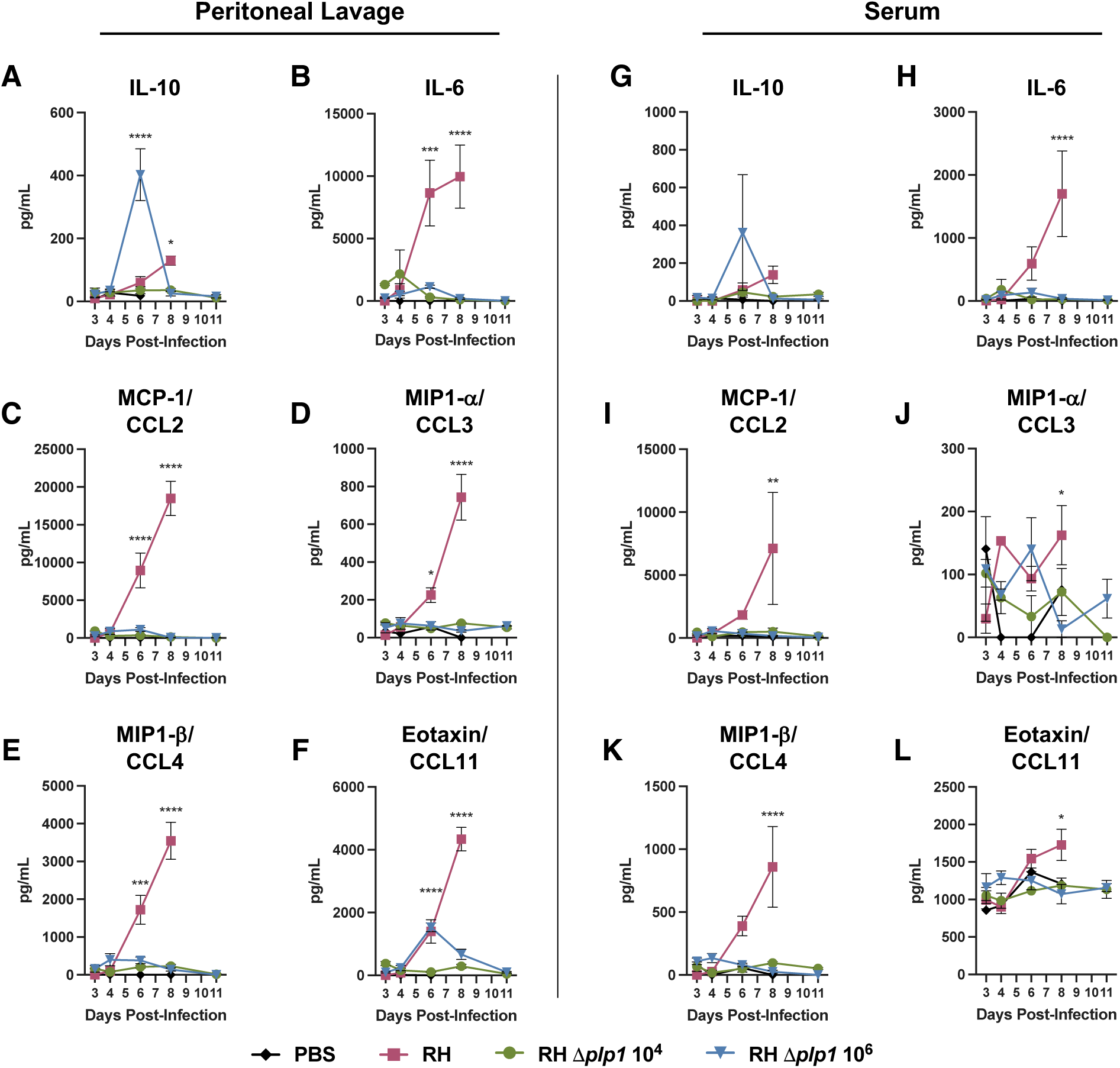
Dysregulated inflammatory cytokine responses characterize wild type infection compared with PLP1 deficient parasite infection. PEC lavage (2 mL PBS wash of the peritoneal cavity, A-F) and serum (G-L) cytokine and chemokine responses were measured after i.p. infection of C57BL/6 mice with either 10^2^ WT (RH, red), 10^4^ RH *Δplp1* (green), or 10^6^ RH *Δplp1* (blue) tachyzoites. IL-10 (A, G), IL-6 (B, H), MCP-1/CCL2 (C, I), MIP1-a/CCL3 (D, J), MIP1-b/CCL4 (E, K), and Eotaxin/CCL11 (F, L) were assessed by Milliplex between 3 and 11 dpi as described. Graphs are representative results from one of two experiments (n=3 with a single PBS control per day). Two-way ANOVA statistical analysis was used to determine significance; * represents p ≤ 0.05, ** represents p ≤ 0.01, *** represents p ≤ 0.001. All WT infected mice succumbed by days 8 and 9 post-infection.

## Discussion

The work presented here provides a first insight into the role of the *Toxoplasma* pore forming toxin PLP1 in the pathogenesis of toxoplasmosis in mice. Prior work from our laboratory showed that although PLP1 is required for efficient egress from the parasitophorous vacuole, PLP1 deficient parasites can be propagated normally in cell culture (26–29). The dispensability of PLP1 in cultured parasites is also supported by *PLP1* having a positive phenotype score from a genome-wide CRISPR screen that evaluated the level to which individual genes conferred fitness during *in vitro* replication (44). However, in results repeated here in C57BL/6 mice, *PLP1* knockout parasites are avirulent despite their ability to replicate *in vivo.* This differs significantly from the parental RH strain, where a single parasite is acutely virulent (9, 15, 27). The data presented here suggest that attenuated virulence of *PLP1* knockout parasites is associated with both a control of parasite replication and the regulation of the host immune response during the acute phase of infection. In the present work we interrogated several plausible hypotheses to understand how PLP1 expression confers a virulence enhancing phenotype during infection by the parental RH strain.

Pore forming toxins (PFTs), including MACPF proteins, are virulence factors expressed by multiple bacterial pathogens, including *Listeria monocytogenes*, *Streptococcus pyogenes*, and *Bacillus anthracis* (45, 46). PFTs promote virulence in multiple ways including increasing pathogen survival (*i.e*., *Listeria* and *Mycobacterium* escape from the phagosome via listeriolysin O (LLO) and ESAT-6, respectively), activation of proinflammatory responses (*i.e*., MAPK pathway activation by pneumolysin and streptolysin O (SLO), or inflammasome activation by LLO and SLO), and immune evasion through immune cell death and manipulation of host responses (46, 47). Because the mechanism by which *Toxoplasma’s* PLP1 influences parasite virulence remains unknown, we initially assessed whether parasites lacking PLP1 can infect and disseminate throughout an infected mouse. Indeed, our data suggest that *Δplp1* parasites replicate after injection and establish a multisystemic infection, but replication is eventually controlled and parasite burden resolved. We previously showed that *Δplp1* parasites are characterized by an egress defect, so it is plausible that these parasites are still able to employ a Trojan Horse mechanism by invading dendritic cells and thus disseminating throughout the organism (48–50).

Infection with wild type RH parasites results in a robust type 1 immune response that is lethal in mice (16, 17). Since our findings suggest that *Δplp1* parasites establish an acute infection, we next investigated if host immune response differences may explain the observed attenuation in the mouse virulence phenotype. Our data showed that i.p. infection with WT parasites resulted in a widespread necrosis and profound depletion of monocytic and dendritic peritoneal cells, that was not observed during infection with *Δplp1* parasites, regardless of the infection inoculum used. The impact of bacterial PFTs on the depletion of relevant immune cells may offer insight into the mechanism of *Toxoplasma’s* PLP1 activitiy. Examples for these mechanisms include *Listeria’s* LLO mediating phagosome evasion and activating the NLRP3 inflammasome, and *Staphylococcus aureus’s* leucocidin ED or *S. pyogenes* SLO and streptolysin S (SLS) critical role in the elimination of host phagocytes (47, 51, 52). Future studies will investigate whether monocyte and dendritic cell depletion is the result of direct PLP1-mediated lysis during parasite egress, or if it is a defect in hematopoiesis, including lymphomyeloid differentiation, or by activating a programmed cell death pathway such as the inflammasome, active cytotoxicity, or a currently unknown immune process.

The depletion of dendritic cells observed in WT infected mice in our current study substantiated previous reports that described decreased dendritic cell numbers as a characteristic of virulent RH infection, but not avirulent Type II infection (36). The authors further described RH infection-specific impairment of dendritic cell early activation compared with Type II infection (36). Similar to Type II infection, dendritic cells responding to avirulent *Δplp1* parasites were phenotypically more activated at 3 dpi than those in RH infected mice, as measured by CD86 expression. Likewise, infiltrating Ly6C^+^ monocytes were also more activated 3 days post *Δplp1* infection compared to RH infection. The impact of early antigen presenting cell activation during an avirulent infection is not fully understood, but Tait *et al.* demonstrate that downstream effects on the CD8^+^ T cell responses are partially responsible for controlling parasite burden (36). Considering these results together, the *Δplp1* isogenic RH strain infers a role for PLP1 as a virulence factor that promotes *Toxoplasma’s* ability to alter myeloid cell populations, and specifically antigen presenting cell activation.

Like other PFTs, our data show that PLP1 contributes to recruitment of neutrophils to the site of infection coincident with increased production of G-CSF, CXCL2 and CXCL1 chemokines, and consistent with previous observations that signalling through the receptor for these chemokines is important for neutrophil migration to the site of infection (53). Strikingly, when we compare infection with WT and *Δplp1* parasites, chemokine levels rise significantly above naïve in the infection with WT parasites. These results are similar to those observed in the *S. aureus* Panton-Valentine leukocidin (PVL) model where PVL-induced inflammation results in the influx of more neutrophils through increased production of chemoattractants like CXCL1 (KC) (54). Inflammation and bacterial virulence were attenuated when PVL was antibody neutralized or when mice were infected with PVL deficient *S. aureus* (54), analogous to our observations during infection with PLP1*-*deficient *T. gondii*.

It was previously reported that neutrophils do not contribute significantly to *T. gondii* infection outcomes, but these studies were conducted using the avirulent, cystogenic Prugniaud strain (Type II) (35). To date, no studies have specifically investigated the role of neutrophils beyond the first 36 hours of an RH infection (37, 38, 53). This study shows that WT RH infection, but not *Δplp1* infection, significantly increases both chemoattractants and the growth factor G-CSF, driving significant increases in neutrophil numbers and influx to the peritoneal cavity. Our data establish that prior to mice succumbing to WT infection, neutrophils are the dominant cell type observed in the peritoneal cavity. In view of our results, neutrophils appear to contribute to RH virulence by inducing dysregulated bystander immunopathology (39). Attenuated neutrophil recruitment coupled with the prevailing population of lymphomyeloid cells at the site of infection may be, at least in part, responsible for the decreased virulence of the *Δplp1* strain.

Pathology during acute toxoplasmosis has been linked to an overstimulation of the host immune system (16, 17). Like these other studies, evidence for this phenomenon in our study included unregulated serum IFN-γ and IL12-p40 production after infection with WT parasites compared to *Δplp1* infected mice. Our neutrophil, cytokine, and chemokine data are also consistent with the concept that RH virulence is associated with an overstimulated immune response. Additionally mice infected with WT parasites showed local and systemic spikes in IL-10, TNF-α, IL-18, IL-6, CCL2, CCL3, CCL4, CCL11, G-CSF, and CXCL10 just prior to succumbing to infection, consistent with prior observations (17) and indicative of an agonal overinduction of proinflammatory cytokine activity (55, 56). Significantly, many of the upregulated cytokines and chemokines found in response to WT infection, are among those that typify a “cytokine storm” responsible for adverse outcomes in viral and bacterial infections, immunotherapies, cancers, and autoimmune diseases (57). Studies have demonstrated that PFTs play an important role in the early activation of proinflammatory pathways leading to increased cytokines like IL-1β, TNF-α, and IL-6 (46). Infection with PLP1-deficient parasites resulted in drastically reduced TNF-α and IL-6, resolved infection-driven IFN-γ production, and failed to result in exaggerated cytokine and chemokine responses associated with proinflammatory overproduction and host mortality (17, 43). The mechanism for PFT associated proinflammatory activation is unclear, though it is known that PFTs are involved in the activation of host responses through mechanisms like activation of MAPK pathways, pattern recognition receptor ligation, and inflammasome activation (46). Additionally, PFTs have demonstrated roles in direct manipulation of host proteins; as with *Streptococcus pneumoniae*’s pneumolysin activation of the host deubiquitinating enzyme CLYD, or the delivery of bacterial virulence proteins into the cell cytoplasm like *Bacillus anthracis’s* delivery of lethal factor by protective antigen (46). Further studies are needed to fully probe the extent to which PLP1 impacts cellular danger recognition pathways and promotes the proinflammatory environment observed during infection. In summary, we propose that PLP1, or factor(s) associated with the function of PLPI, is a potent immune and inflammatory mediator, as observed with PFTs in other pathogens.

## Materials & Methods

### Parasite strain generation

The RH *Δ*KU80 *Δ*HPT GFP Luciferase strain was generated by co-transfecting RH *Δ*KU80 *Δ*HPT purchased from ATCC (PRA-319) with 5µg of pTOXO_Cas9_CRISPR plasmid (58) expressing a UPRT guide – 5’ GGCGTCTCGATTGTGAGAGC - and 3µg of the pGRA1-GFP-pDHFR-Luciferase repair construct. Transgenic parasites were selected with floxuridine (FUDR). Stable clones with similar luminescence activity were selected for intra-vital imaging as previously described (59). *Plp1* knockouts were engineered in this strain by insertion of HXGPRT into the genomic locus using Cas9-CRISPR and 2 *plp1* guides – 5’ GTGTGGACCATGGAATGCTG and 5’ GTCTCTCCAGCGCTGTCAAA (5µg per plasmid), and confirmed by locus sequencing. All repair constructs were generated by Roche KAPA Taq mediated polymerase chain reaction and consisted of 30 base pair homologous ends.

### Host cell and parasite culture

All cells and parasites were maintained in a humidified incubator at 37°C and 5% CO_2_. Human foreskin fibroblast cells (HFF, ATCC CRL-1634) were maintained in Dulbecco’s modified Eagle’s medium (DMEM) supplemented with 10% fetal bovine serum, 20 mM HEPES pH 7.4, 2 mM L-glutamine, and 50 µg/mL penicillin/streptomycin. All *T. gondii* parasites were maintained by serial passaging in HFF cells and were routinely checked for mycoplasma contamination. In preparation for infection, freshly lysed parasites were filtered through a 3 µm Nuclepore Track-Etch membrane (Whatman, Cytiva), washed once with phenol free RPMI Medium (Gibco, ThermoFisher), and resuspended in 1x PBS (Gibco, ThermoFisher).

### Mice

Female 6-8 week old C57BL/6 mice (strain 000664, Jackson Laboratories, Bar Harbor, ME) were purchased and maintained under specific pathogen free conditions in the National Institutes of Health/National Institutes of Allergy and Infectious Disease (NIAID) Animal Facilities for the study. All procedures were approved by the NIAID Animal Care and Use Committee (ACUC), Animal Study Protocol LPD-22E, in accordance with the American Association for the Accreditation of Laboratory Animal Care (AAALAC).

### Bioluminescence imaging

Three to four C57BL/6 mice were infected with either 10^2^ (RH), 10^4^ (*Δplp1*), or 10^6^ (*Δplp1*) freshly lysed tachyzoites in 200 µL PBS via intraperitoneal (ip) injection. Mice were bioluminescence imaged (BLI) at 24 hours, and 2, 4, 6, 8, and 11 days thereafter. To do this, mice were intraperitoneal (ip) injected with 3 milligrams luciferin (in 200 µL PBS), maintained for 5 minutes for substrate dissemination and then anesthetized in an oxygen rich chamber with isoflurane. Mice were imaged using an IVIS BLI system (PerkinElmer, Caliper Life Sciences) in ventral and lateral recumbency. For luminescent imaging of organs, mice were i.p. injected with luciferin as described, humanely euthanized after 5 minutes, and the organs placed in warm phenol red free RPMI medium and immediately imaged (**Fig 2**). Two milliliters of PBS were used to rinse the peritoneal cavity (PL) and placed into 20 mm dishes for imaging with the excised organs (**Fig 2**). Parasite burden in whole mice and specific organs was measured as the total flux (photons/second) within a region of interest (ROI) as analyzed with Aura Imaging Software (Spectral Instruments Imaging).

### Infection analysis

C57BL/6 mice were infected with either 10^2^ wild type (RH), 10^4^ Δ*plp1*, 10^5^ Δ*plp1* or 10^6^ Δ*plp1* parasites, as described above. At 3, 4, 6, 8, and 11 days post-infection (dpi) mice were humanely euthanized, serum was collected by cardiac puncture, and organs were collected for luminescent imaging (see above) and histopathology. Samples from the liver and spleen of each mouse in the challenge cohort and control were collected and preserved in Hollande’s fixative (StatLab). Tissues were processed by conventional histologic techniques, embedded in paraffin, sectioned at 5 µm and stained with hematoxylin and eosin. The prepared slides were blinded and reviewed. Lesions and any discernible protozoa were initially identified, qualitatively scored, and the data tabulated. Histopathology was scored based on lesion intensity and distribution, then the findings were calibrated for the 3-4 tissues present on each examined slide.

After BLI imaging of the organs, peritoneal exudate lavage was immediately retrieved from the 20 mm dishes (see above) for further analysis. Lavage was centrifuged (1250 rpm, room temperature) for 5 minutes and peritoneal exudate cells (PECs) and supernatant were collected for further flow cytometry and cytokine/chemokine analysis, respectively. A subset of the infected mice were maintained for the course of infection to collect weight and survival data.

### Cytokine and chemokine analysis

Cytokine concentrations in both the serum and the peritoneal lavage were determined by IL-12p40 (Cat. # DY499 Duosets, R&D Systems, Minneapolis MN) and IL-18 ELISA (Cat. # BMS618-3, ThermoFisher Scientific) using the manufacturer’s recommended protocol, modified to include an overnight sample incubation. Additionally, a panel of 20 cytokines/chemokines were assessed using a custom MilliporeSigma Milliplex kit (EMD Millipore Corporation, Burlington MA); analytes included were IFN-gamma, IL-10, IL-12p70, IL-15, IL-1alpha, IL-2, IL-6, IP-10/CXCL10, KC/GRO/CINC-1/CXCL1, MCP-1/CCL2, MIP-1alpha/CCL3, MIP-1beta/CCL4, MIP-2/CXCL2, RANTES/CCL5, TNF-alpha, Eotaxin/CCL11, LIX, MIG/CXCL9, G-CSF, and GM-CSF. Milliplex analytes were read on a BioMax.

### Flow cytometry

PECs from infected C57BL/6 mice were centrifuged at 1,200 rpm for 5 minutes, and washed with PBS before staining with Invitrogen LIVE/DEAD Fixable Blue stain (UV450, ThermoFisher Scientific). Cells were washed and blocked (eBioscience CD16/CD32 clone 93, FACs buffer - 0.2% BSA in PBS), followed by incubation with Ly6G (PE, 1A8, BD Pharmingen), CD11c (PE-CF594, N418, BD Biosciences), CD24 (PE-Cy7, M1/69, BioLegend), CD86 (APC, GL-1, Biolegend), CD45 (Alexa Fluor 700, 30-F11, BioLegend), F4/80 (APC-Cy7, BM8, BioLegend), MHCII (Brilliant Violet 421, M5/114.15.2, BioLegend), CD11b (Brilliant Violet 605, M1/70, BioLegend), and Ly6C (Brilliant Violet 785, HK1.4, BioLegend). The FITC channel was left open for parasite specific GFP. Finally, the cells were fixed with 4% PFA and left in FACs Buffer until analysis. Data was acquired on an LSRII Fortessa with FACs Diva software, and later analyzed using FlowJo software (BD Biosciences). Gating strategy provided (**S1 Fig**). CD86 offset histogram plots were generated using FlowJo.

### Statistical analysis

Graphs and statistical analyses were performed using GraphPad’s Prism software, version 9.3.1. All data was analyzed for statistical significance by one-way or two-way analysis of variance (ANOVA) following Tukey’s multiple comparisons test model. Significance was determined as *=p≤0.05, **=p≤0.005, ***=p≤0.0005, and ****=p≤0.0001.

## Acknowledgements

We gratefully acknowledge the funding support of the U.S. National Institutes of Health (R01AI46675 to VBC). This work was supported in part by the Division of Intramural Research (DIR) of the National Institute of Allergy and Infectious Diseases (NIAID) at the National Institutes of Health (MEG). The funders had no role in study design, data collection, data analysis, publishing decisions, or manuscript preparation and the opinions expressed in this article are the author’s own. This research was conducted prior to Dr. Guerra’s employment at the NIH. We also acknowledge Jay Bream, Ph.D. at the Johns Hopkins Bloomberg School of Public Health for insightful comments on data interpretation and presentation.

## Author contributions and notes

Beth Gregg: Conceptualization, Data curation, Formal analysis, Investigation, Methodology, Writing – review & editing

Alfredo J. Guerra: Writing – original draft, Writing – review & editing

Stephen A. Raverty-Formal analysis, Writing – review & editing

Aline Sardinha-Silva: Investigation, Methodology

Bjorn F.C. Kafsack: Conceptualization, Data curation, Formal analysis, Investigation, Methodology, Writing – review & editing

Tracey L. Schultz: Investigation, Methodology

Stephen J. Gurczynski: Conceptualization, Data curation, Writing – review

Bethany B. Moore: Conceptualization, Writing – review

Michael E. Grigg: Conceptualization, funding acquisition, Writing – review & editing

Vern B. Carruthers: Conceptualization, funding acquisition, Writing – review & editing

The authors declare no conflict of interest.

## Supporting Information

**S1 Fig.**
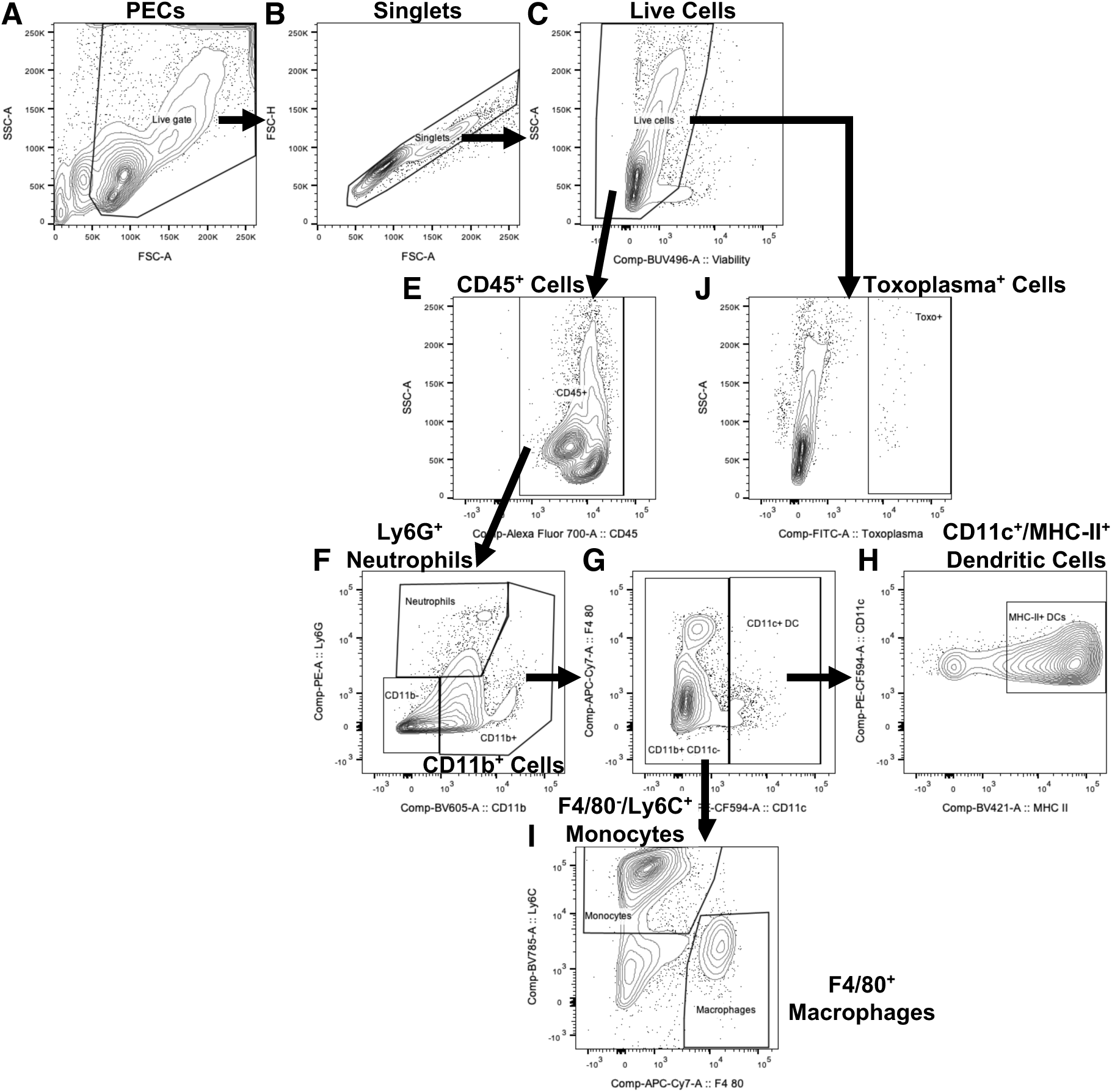
Gating strategy for flow cytometric analysis. PEC immune cell populations referenced in figures 4 and 5, were identified based on sequential phenotyping by immunological markers. PECs were gated based on FSC-A/SSC-A (A) followed by doublet exclusion (B - singlets, FSC-A/FSC-H). The live cell population was delineated by low levels of LiveDead Fixable Blue staining (C) and hematopoetic cells were gated as CD45^+^ (E). From the CD45^+^ cells, neutrophils were gated as CD11b^−^/Ly6G^+^ (F) and the CD11b^+^ cell population was further divided by expression of CD11c (G). CD11c^+^ cells were confirmed as dendritic cells by expression of MHC-II (H), while CD11c^−^ cells were phenotyped as either F4/80^+^ macrophages or F4/80^−^/Ly6C^+^ monocytes (I). Total *Toxoplasma* infected PECs were accounted for by GFP expression in the CD45^+^ cell population (J). Diagram is a representative example from the WT RH infected mice 4 dpi.

**S2 Fig.**
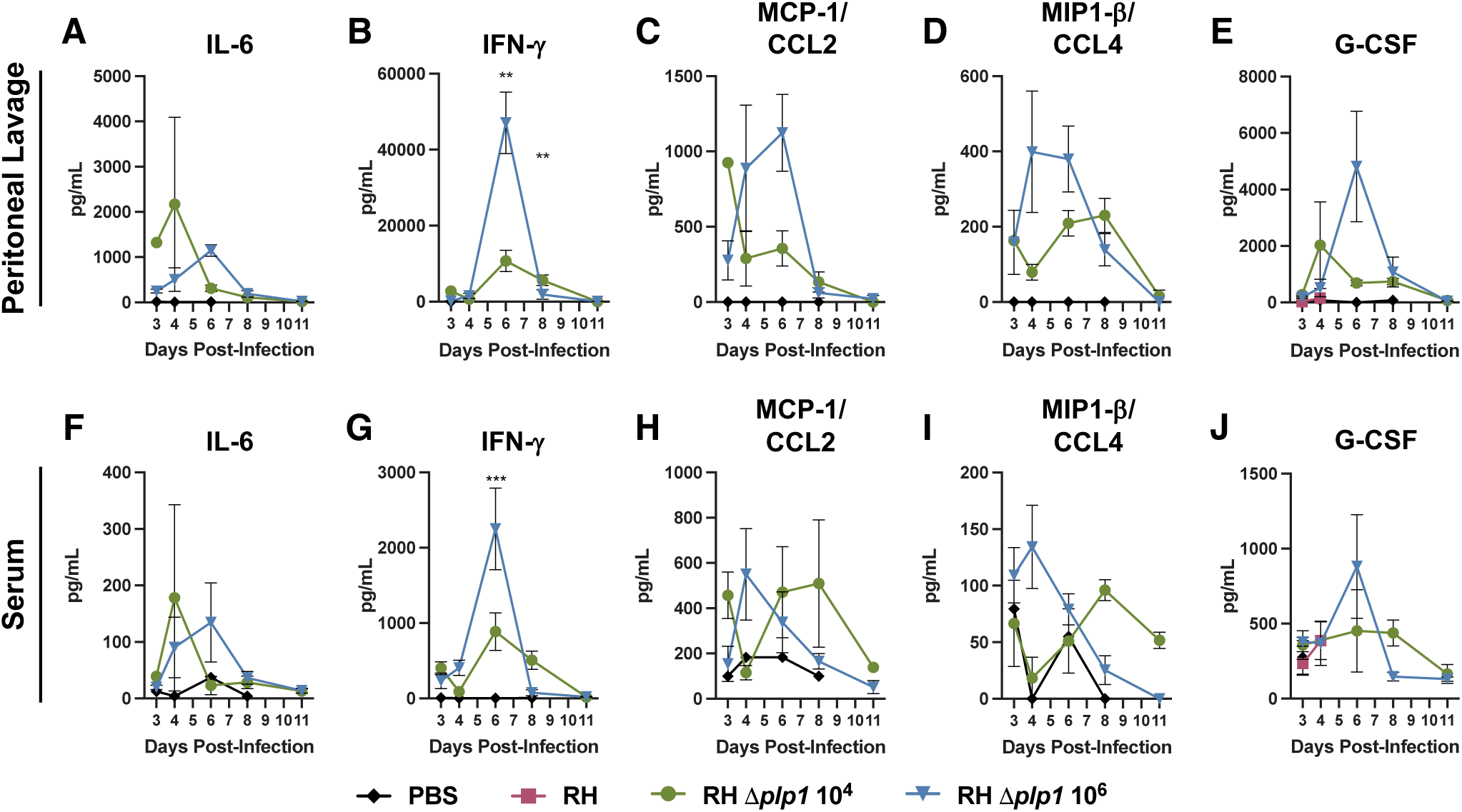
RH PLP1 knockout parasites induce cytokine and chemokine responses after infection. PEC lavage (A-E) and serum (F-J) cytokine and chemokine responses were measured and graphed for 10^4^ *Δplp1* (green), or 10^6^ *Δplp1* (blue) i.p. infection of C57BL/6 mice. IL-6 (A, F), IFN-g (B, G), MCP-1/CCL2 (C, H), MIP1-b/CCL4 (D, I), and G-CSF (E, J), were assessed by Milliplex between days 3 and 11 post-infection, as described. Graphs are representative results from one of two experiments (n=3 with a single PBS control per day). Data for RH infection is omitted to better demonstrate changes induced by PLP1 deficient parasite infection. Two-way ANOVA statistical analysis was used to determine significance; ** represents p ≤ 0.01, *** represents p ≤ 0.001.

